# Discovery of a microbially produced small molecule in a host-specific organ

**DOI:** 10.1101/2020.08.17.254854

**Authors:** Katherine E. Zink, Denise A. Tarnowski, Phillip R. Lazzara, Terry W. Moore, Mark J. Mandel, Laura M. Sanchez

**Affiliations:** Department of Pharmaceutical Sciences, University of Illinois at Chicago, Chicago, IL 60612; Department of Medical Microbiology and Immunology, University of Wisconsin - Madison, Madison, WI 53706; University of Illinois Cancer Center, University of Illinois at Chicago, Chicago, IL 60612

## Abstract

The lifelong relationship between the Hawaiian bobtail squid, *Euprymna scolopes*, and its microbial symbiont, *Vibrio fischeri*, represents a simplified model system for studying microbiome establishment and maintenance. The bacteria colonize a dedicated symbiotic light organ in the squid, from which bacterial luminescence camouflages the hosts in a process termed counterillumination. The squid hosts hatch without their symbionts, which must be acquired from the ocean amid a diversity of non-beneficial bacteria, so precise molecular communication is required for initiation of the specific relationship. It is therefore likely that there may be specialized metabolites used in the light organ microenvironment to modulate these processes. To identify small molecules that may influence the establishment of this symbiosis, we used imaging mass spectrometry to analyze metabolite production in *V. fischeri* with altered biofilm production, which correlates directly to colonization capability in its host. ‘Biofilm-Up’ and ‘Biofilm-Down’ mutants were compared to a wild-type strain, and masses that were more abundantly produced by the biofilm-up mutant were detected. Using a combination of structure elucidation and synthetic chemistry, one such signal was determined to be a diketopiperazine, cyclo(d-histidyl-l-proline). This diketopiperazine modulated luminescence in *V. fischeri* and, using label-free imaging mass spectrometry, was directly detected in the light organ of the colonized host. This work highlights the continued need for untargeted discovery efforts in host-microbe interactions and showcases the benefits of the squid-*Vibrio* system for identification and characterization of small molecules that modulate microbiome behaviors.

**Significance Statement:** The complexity of animal microbiomes presents challenges to defining signaling molecules within the microbial consortium and between the microbes and the host. By focusing on the binary symbiosis between *Vibrio fischeri* and *Euprymna scolopes*, we have combined genetic analysis with direct imaging to define and study small molecules in the intact symbiosis. We have detected and characterized a diketopiperazine produced by strong biofilm-forming *V. fischeri* strains that was detectable in the host symbiotic organ, and which influences bacterial luminescence. Biofilm formation and luminescence are critical for initiation and maintenance of the association, respectively, suggesting that the compound may link early and later development stages, providing further evidence that multiple small molecules are important in establishing these beneficial relationships.

## Introduction

Nearly all animals are hosts to microbial species.(1) Communities of microbes can inhabit host organs intracellularly or extracellularly, may confer benefits to the host, and collectively are referred to as the microbiome.(2) Some of these relationships have been explored by investigating the role that chemical communication plays between species, whether to initiate interactions or to maintain them.(3) For instance, the plant pathogen, *Ralstonia solanacearum* produced a hybrid non-ribosomal peptide synthetase-polyketide synthase (NRPS-PKS) lipopeptide that induced chlamydospore development in soil dwelling filamentous fungi which may help it persist in the soil environment. Whereas the fungal species *Fusarium fujikuroi* and *Botrytis cinerea* produce bikaverin to inhibit invasion of ralsolamycin producing strains of *R. solanacearum*.(4, 5) In cheese rind-derived microbial interactions, small molecules like Zinc-coproporphyrin III and other siderophores mediate trace metal acquisition between bacteria and fungi.(6, 7) In both of these cases, specialized metabolites were central to understanding the observed phenotypes between the domains of life. Beyond antagonistic relationships, in many bacterial species, the density of the population is monitored by self-secreted “autoinducer” compounds, which includes homoserine lactones in Gram-negative bacteria and small peptides in many Gram-positive organisms.(8) Quorum sensing was discovered in *Vibrio fischeri*, a marine Gram-negative bacterium that colonizes squid and fish hosts. Studying *V. fischeri* in the context of its natural hosts provides an opportunity to expand our knowledge of biologically relevant compounds that influence microbial behaviors and microbe-host signaling in animal microbiomes.

To study the role of specialized metabolites in microbiome-host interactions, we have focused on the binary partnership between *V. fischeri* and the Hawaiian bobtail squid, *Euprymna scolopes*. Shortly after hatching, the squid host acquires the symbiont from the seawater, and the microbe proceeds to colonize the epithelial-lined crypts of the light organ. The bacteria gain a safe nutrient-rich habitat in which they can replicate.(9) In turn, the bacteria in the aptly-named “light organ” produce luminescence for the squid. The bacterial light camouflages the nocturnal predator via counterillumination, hiding the animal’s shadow in the moonlight so that it is less visible to predators and prey while the squid forages.(10) Many aspects of the colonization process are comparable to bacterial colonization of human epithelial cells in the gut and on skin.(3, 11, 12) As a naturally co-evolved relationship where one host organ houses only one bacterial species, the simplified *Vibrio*-squid system allows for analysis of specialized metabolites involved in colonization of epithelial tissues. The colonization process is highly efficient and the host tissues are largely transparent, facilitating imaging approaches during the early stages of colonization. *V. fischeri* is amenable to genetic manipulation, including mimicking of symbiotic behaviors in culture. Furthermore, both partners can be raised separately and then mixed in a controlled fashion, enabling well-controlled experiments.

There is remarkable colonization specificity in that only *V. fischeri*--and only specific strains of *V. fischeri--colonize* the *E. scolopes* light organ.(13) Part of this specificity lies in the exchange of chemical cues between the two partners that leads to maturation of the symbiosis, including the release of bacterial products peptidoglycan, the peptidoglycan monomer tracheal cytotoxin (TCT), lipopolysaccharide (LPS); and host release of nitric oxide and chitin.(14–19) A major checkpoint for this specificity is that bacterial biofilm production is required for bacterial aggregation and subsequent colonization.(20, 21) Strains lacking the biofilm regulator RscS (specifically ES114 *rscS* rscS*::Tn*erm*, in this study) do not synthesize the symbiotic biofilm and are unable to robustly colonize the squid host.(22) Wild-type ES114 does not form biofilm in typical lab culture conditions but does form symbiotic biofilm aggregates in the squid host.(20, 23, 24) Additionally, a strain carrying both upregulation of the biofilm activator *rscS* and a deletion in the biofilm inhibitor *binK* activates formation of a substantial amount of symbiotic biofilm, which is evident as wrinkled colony biofilms in culture (ES114 *rscS* ΔbinK*).(25) Therefore, in this study we used this panel of strains to identify compounds that correlated with symbiotic biofilm formation, and we termed them Biofilm-Down, wild type (WT), and Biofilm-Up, respectively. We hypothesized that small molecule production may contribute to the biofilm effects in the host and to the propensity for *V. fischeri* to outcompete other bacteria, and we used these defined strains with cutting edge analytical technologies to address these questions.

One method that can rapidly determine differences in metabolite production of microbes is agar-based imaging mass spectrometry (IMS). This approach has been applied to decipher microbe-microbe chemical communication and changes in metabolite production between microbial colonies grown in different environmental conditions.(6, 26, 27) IMS is especially valuable for the comparison of metabolites of defined strains because changes in production, as measured by ion intensity, can be specifically and rapidly detected and evaluated in the context of genetic differences.(28) IMS analysis of WT *V. fischeri* compared to Biofilm-Up and Biofilm-Down strains revealed that several metabolites were significantly upregulated in Biofilm-Up compared to the other strains. These metabolites were hypothesized to be important for the production of the symbiotic biofilm and/or the increased fitness of the Biofilm-Up mutant in the light organ environment. Additionally, because of the small size of *E. scolopes* hatchlings and the anatomical accessibility of the light organ, a whole-body imaging approach was optimized for *in vivo* investigation of the light organ.(29) Together, these two IMS approaches were utilized to determine differences in specialized metabolism in *V. fischeri mutants*, both *in vitro* and *in vivo* and led to the isolation and structure elucidation of a small molecule that increases *V. fischeri* luminescence, which may provide a fitness advantage in host-microbe recognition.

## Results and Discussion

### *Untargeted spatial metabolomics of* V. fischeri *biofilm mutants*

IMS was employed to generate a rapid screen of mass-to-charge ratios (*m/z*’s) in WT, Biofilm-Up, and Biofilm-Down *V. fischeri* to identify ions that differ between solid agar colonies of the three strains. A positive mode analysis in the small molecule range (100-1000 Da) generated a panel of masses that were significantly more abundant (p < 0.1) in the Biofilm-Up mutant than in the other two samples, as determined by the SCiLS software. **Figure S1** displays seven of these significant features that replicated at least twice across four biological replicates. One of these signals, *m/z* 257, was detected with high signal intensity in Biofilm-Up, and low signal intensity in WT and Biofilm-Down (**Figure 1**). This signal was therefore prioritized for dereplication, the process of identifying known unknowns, because biofilm production is strongly correlated to a colonization advantage.

**Figure 1.**
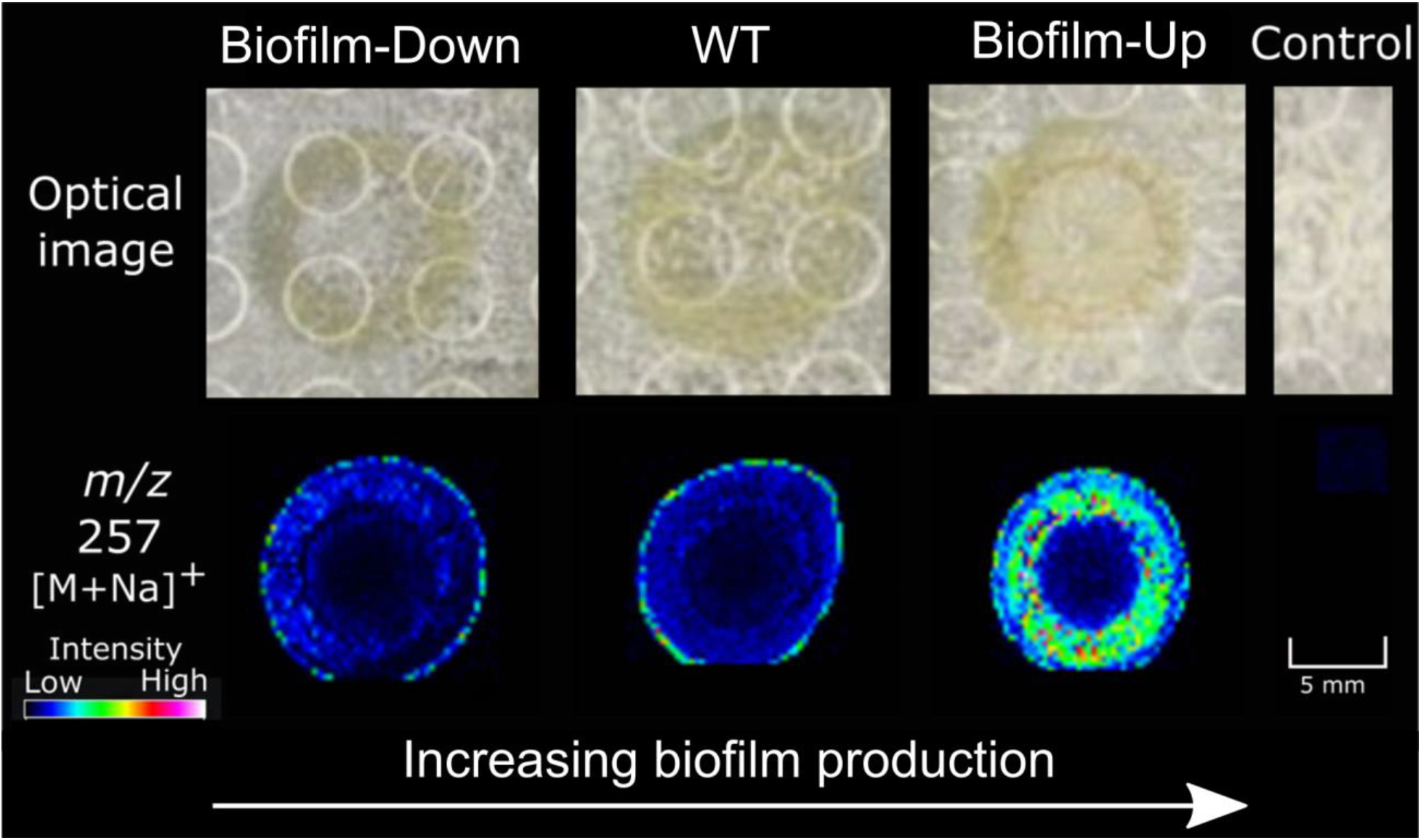
The *m/z* 257 from the mass panel in **Figure S1** was statistically more abundant in Biofilm-Up compared to WT and Biofilm-Down, see Methods for detailed genotypes.

### Dereplication and structure elucidation of cyclo(histidyl-proline)

Masses identified via IMS were prioritized to a select list by evaluating their statistical significance. LC-MS/MS data of crude extracts from each mutant were queried using Global Natural Products Social (GNPS) molecular networking, and a spectrum match from the *V. fischeri* Biofilm-Up extract was detected for a small molecule, cyclo(histidyl-proline), with a molecular weight of 234 g/mol (**Figure S2**).(30, 31) The precursor ion from GNPS (*m/z* 235) matched one of the statistically significant masses from IMS analysis: often, ions detected in IMS are sodiated adducts ([M+Na]^+^) because of salt added to the media to support growth of marine bacteria. Therefore, the *m/z* 257 from **Figure 1** was likely the sodiated adduct of cyclo(His-Pro), and the protonated molecule was *m/z* 235 (**Figure S3**). Of interest to this context, cyclo(His-Pro) is a member of the diketopiperazine (DKP) structural class. DKPs are formed from the cyclization of two amino acids and are prevalent natural products that play a variety of roles in microbial relationships.(32)

While GNPS identified cyclo(His-Pro) as a putative assignment, we employed direct infusion to detect and confirm the fragmentation patterns of the *m/z* 235 from the *V. fischeri ΔbinK* extract and a commercial cyclo(l-His-l-Pro) standard. **Figure 2** depicts a near identical match between the fragmentation patterns of both samples, using direct infusion, which was validated using high-resolution electrospray ionization mass spectrometry (HRESIMS) (**Figure S4**). The protonated precursor ion (*m/z* 235.12) and all high-intensity fragments matched between the standard and extracted samples. These data provided strong evidence to support a Level 2 identification, as defined by the Chemical Analysis Working Group (CAWG), of cyclo(His-Pro) in the *V. fischeri* Biofilm-Up extract.(33)

**Figure 2.**
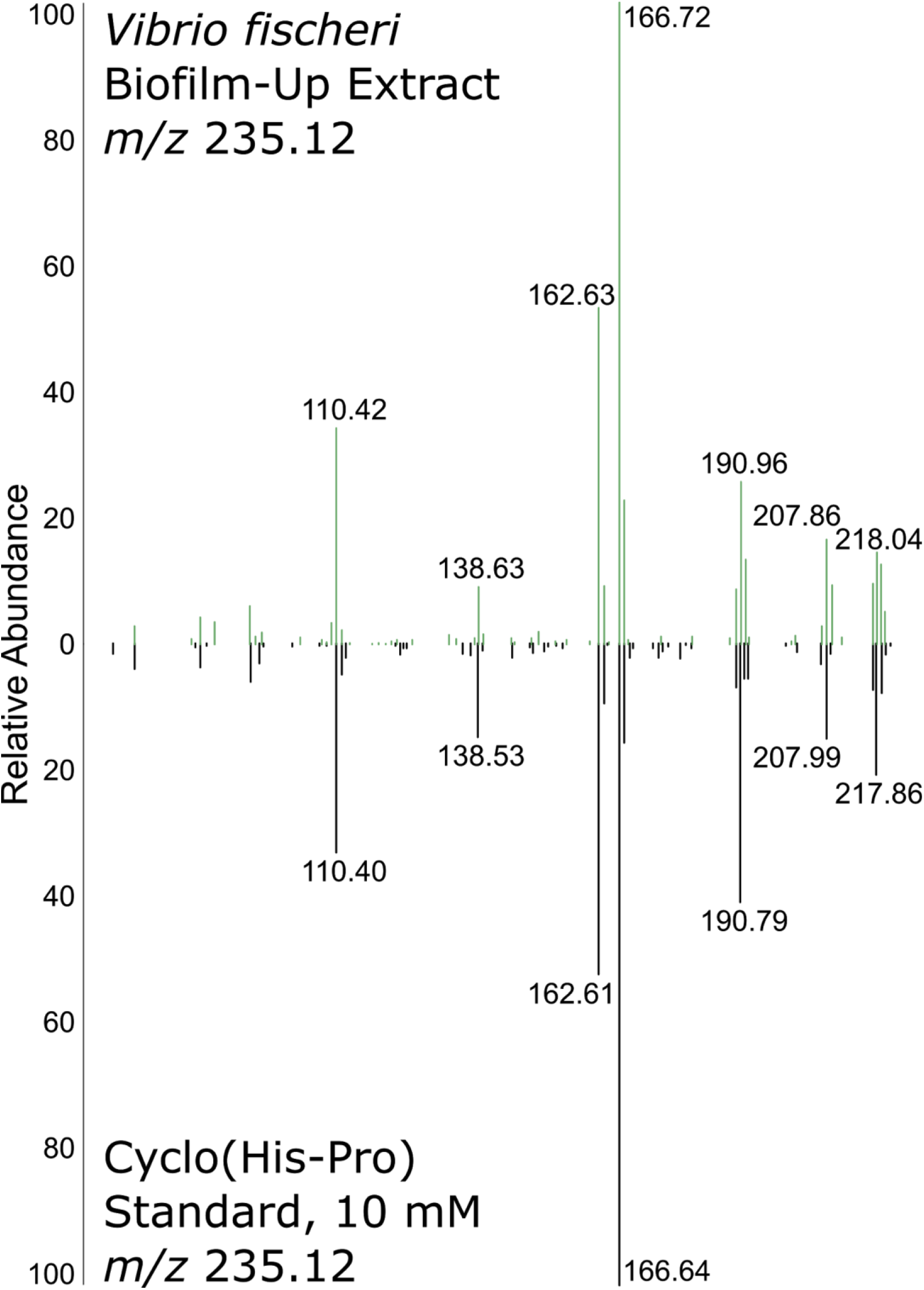
A standard of cyclo(His-Pro) mass fragmentation matched with the extracted molecule at *m/z* 235.12 using direct infusion.

To elucidate the stereochemical configuration of the DKP, as MS/MS is considered to be largely stereochemically blind, we synthesized all four possible stereoisomers of cyclo(His-Pro). Cyclo(l-His-l-Pro) **1**, cyclo(l-His-d-Pro) **2**, cyclo(d-His-l-Pro) **3**, and cyclo(d-His-d-Pro) **4** were synthesized for chemical comparison to the isolated compound from the *V. fischeri* Biofilm-Up extract (**Figure 3A**).(34) To validate the configuration of the synthetic material, each stereoisomer was analyzed using both nuclear magnetic resonance (NMR) and optical rotation (OR). Kukla *et al*. reported optical rotation values, which were used for comparison and validation of each isomer (**SI General Experiment and Compound Characterization and Table S1**).(34)

**Figure 3.**
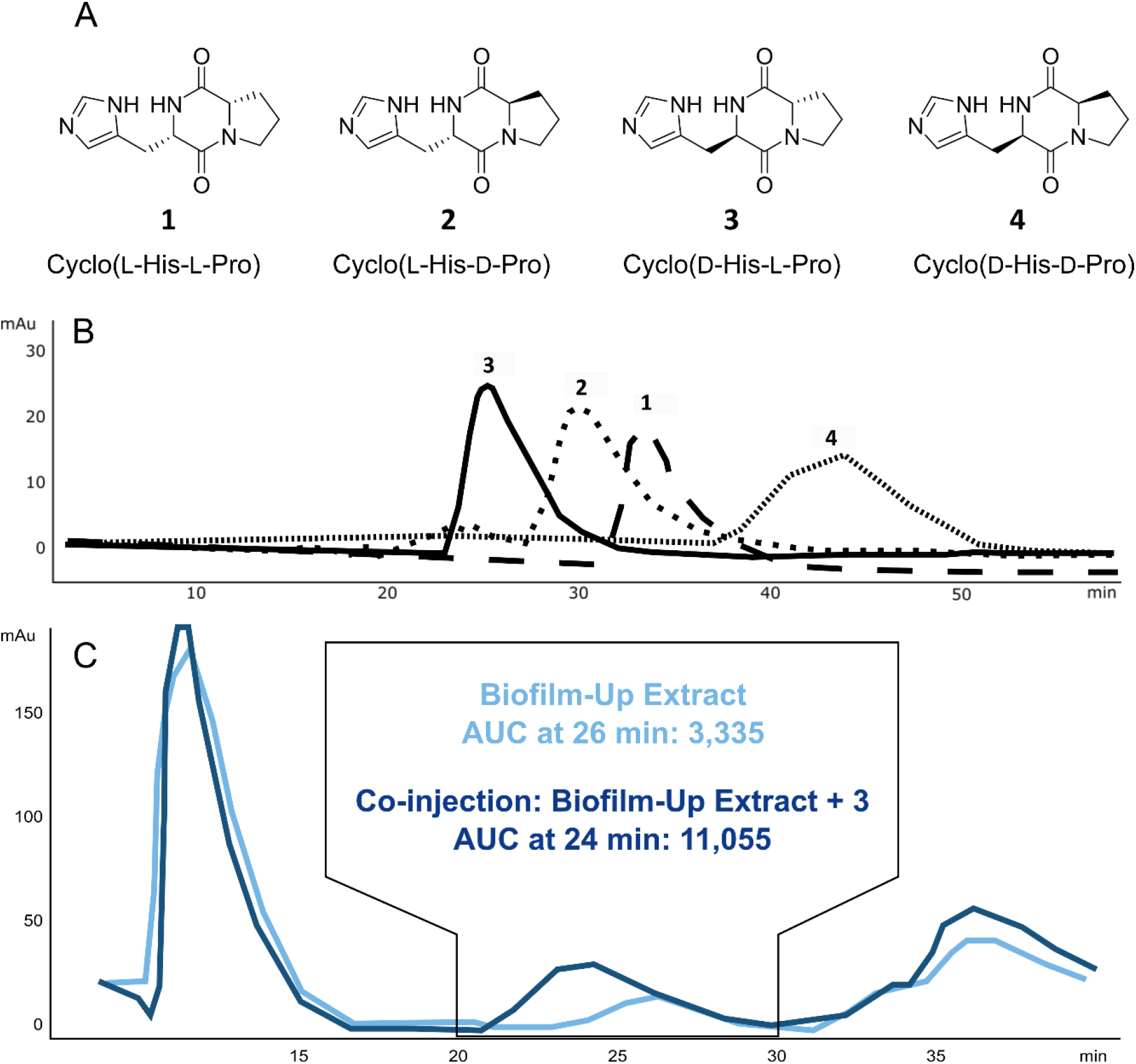
Retention time matching of *V. fischeri* Biofilm-Up extract on a chiral column indicates that the configuration of cyclo(His-Pro) in the microbial extract is stereoisomer **3**, cyclo(d-His,l-Pro). **A)** Structures of all stereoisomers of cyclo(His-Pro). **B)** Retention times of all synthesized stereoisomers: **3** (24 min), **2** (30 min), **1** (33 min), and **4** (44 min). A Cellulose-B column was used to retain stereoisomers using 13:87 IPA:Hexanes over 60 min at 2 mL/min. **C)** A peak at 26 min was observed in the Biofilm-Up extract (light blue trace) and the area under the curve (AUC) measured 3,335. When co-injected with **3** (dark blue trace), the peak at 26 min increased in AUC to 11,055, indicating presence of **3** in the Biofilm-Up extract. UV-vis was monitored at 214 nm.

Chiral chromatography was used to match the retention time of the DKP in the *V. fischeri* extract to all four synthesized stereoisomers (**Figure 3B**). After comparison with each stereoisomer, it was determined that the peak of stereoisomer **3** was also detected in the *V. fischeri* Biofilm-Up extract. When co-injected, the peaks from both samples coalesced to one peak, indicating that their structures and configuration were identical (**Figure 3C**). This retention time matching provides a Level 1 identification for the *m/z* 235 from the *V. fischeri* Biofilm-Up: stereoisomer **3**, cyclo(d-His-l-Pro). The other stereoisomers, namely **2**, were eliminated as being produced in the Biofilm-Up extract because either their retention time did not match an existing peak in the Biofilm-Up extract, or the co-injection peak did not coalesce with an extract peak (**Figure S5**). To attempt to validate the role of **3** in the *E. scolopes* host, we aimed to capture production of **3** *in vivo*.

### In vivo *detection in hatchlings*

A recently developed sample preparation protocol for minimally manipulated whole invertebrate organism IMS was utilized to assess whether **3** was being generated *in vivo* in the light organ of *E. scolopes* hatchlings.(29) The presence of the ion in the light organ would indicate that the DKP has an ecological importance if it can be detected *in vivo*. Three conditions were evaluated: No *V. fischeri* (aposymbiotic), *V. fischeri* WT, and *V. fischeri* WT *ΔbinK* (analogous to the Biofilm-Up strain, but instead of genetically inducing biofilm formation with the *rscS\** allele, relies on induction from native signals in the squid host). *E. scolopes* hatchlings were inoculated with each sample for 3 h, washed, and allowed to continue to colonize for 48 h, as described previously.(35) At 48 h, *m/z* 235 was not detected in the aposymbiotic control and was produced weakly in the WT condition and strongly in the WT *ΔbinK* condition (**Figure 4**). This was evidence that the molecule was produced by the WT strain *in vivo* and that the knockout of *binK* results in increased production of the molecule.

**Figure 4.**
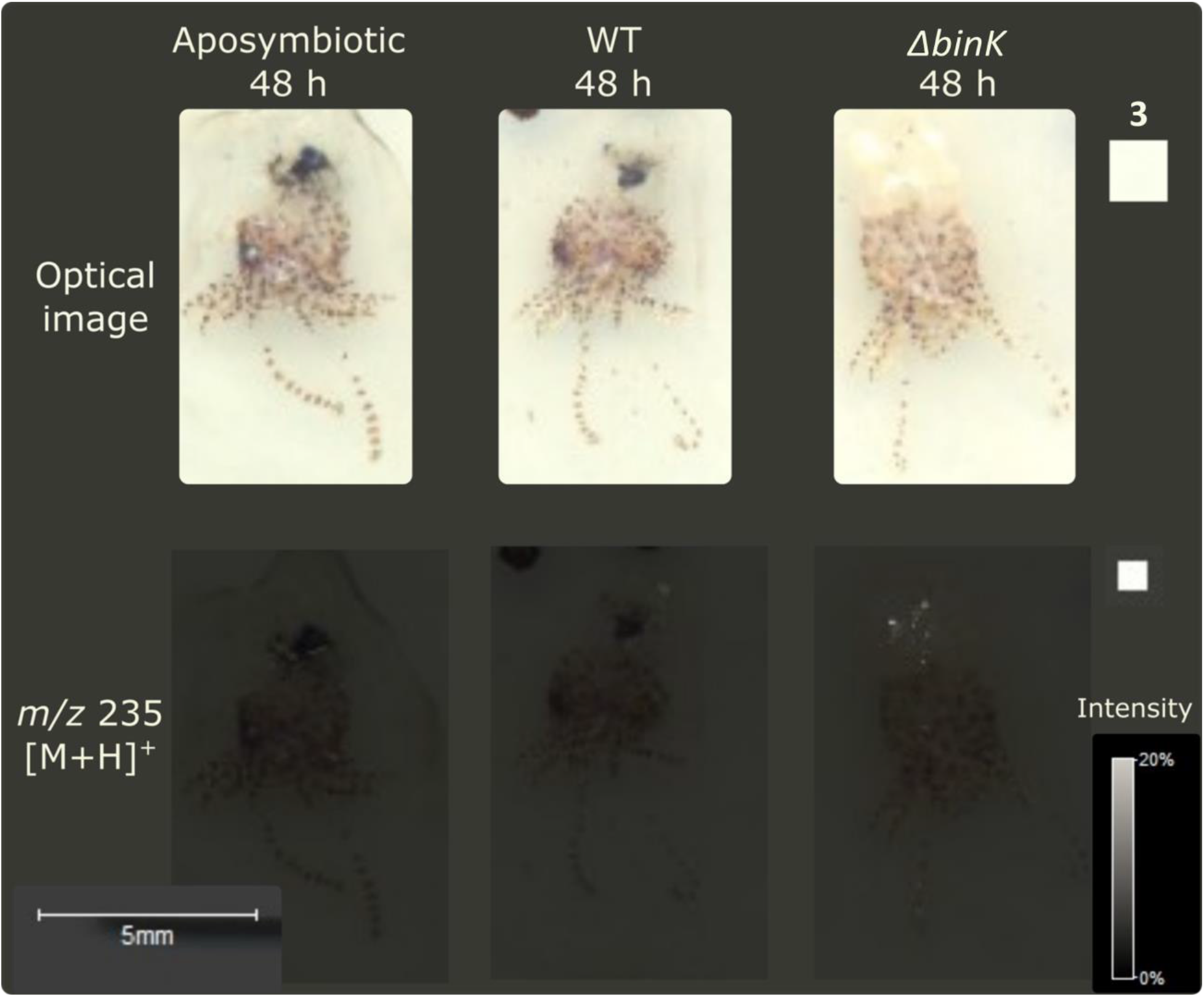
The *m/z* 235 is detected *in vivo* in the light organ of *E. scolopes* hatchlings colonized both when inoculated with *V. fischeri* WT and (strongly) with WT *ΔbinK* (N=3).

The consistency in detection of DKP production *in vitro* and *in vivo* is significant in light of the bacterial strains that were used. The WT strain is the same for both assays, whereas the comparative strain in each case lacks BinK. However, the *in vitro* assay additionally genetically-induces biofilm with an *rscS\** allele (i.e., an overexpressed biofilm regulator), while this allele is not present for the strain used *in vivo*. The rationale for this difference is that the activated RscS allele is used to mimic behavior in culture that is stimulated by the host conditions. Given that DKP production is similarly induced in the host, we conclude that its production is a bona fide output of the symbiotic biofilm pathway and is induced in the host in both WT and in a *ΔbinK* background independent of artificial RscS activation.

### Role in stimulating bioluminescence

Quorum sensing (QS) is the activity of bacterial cells engaging in group behavior that is facilitated by the production of QS molecules by individual cells until a concentration is reached to activate a particular pathway. In the case of *V. fischeri*, the LuxR pathway is activated and bioluminescent activity occurs. The most well studied QS molecules in the *V. fischeri* system are acyl-homoserine lactones (AHL), like N-3-oxohexanoyl-L-homoserine lactone (OHHL). In other bacterial QS systems, proline containing DKPs are responsible for activating the relative pathways. For example, cyclo(l-Pro-l-Leu) produced by *Cronobacter sakazakii*, cyclo(l-Pro-l-Tyr) and cyclo(l-Phe-l-Pro) produced by *Pseudomonas aeruginosa*, and cyclo(l-Pro-l-Val), isolated from *Haloterrigena halophilus* all influence the QS systems of other microbes.(36–38) With so many structural combinations possible, there are many more examples of DKPs that influence microbial QS systems, some in biofilm formation, and some even in *Vibrio spp*.(39–42)

Because of the relationship between biofilm production potential and appearance of **3** in the colonized host, we sought to query whether **3** affected the bioluminescence of *V. fischeri*. An increase in luminescence was observed at low cell densities in the WT strain (ES114) (**Figure 5**). Since ES114 has low luminescence compared to other *V. fischeri* isolates, we tested the effect of the DKP in another strain, EM17, which is a much brighter isolate. This strain also exhibited an increase in luminescence with the addition of exogenous DKP and because it is brighter, the effect is much more pronounced than in ES114 (**Figure 5**). The greatest effect on luminescence in both strains was seen with concentrations of 100 and 250 μM of **3**. The effect of the DKP at low cell densities is similar for other quorum sensing molecules as there is a concentration threshold in these systems, after which the addition of more compound does not increase activity.(43)

**Figure 5.**
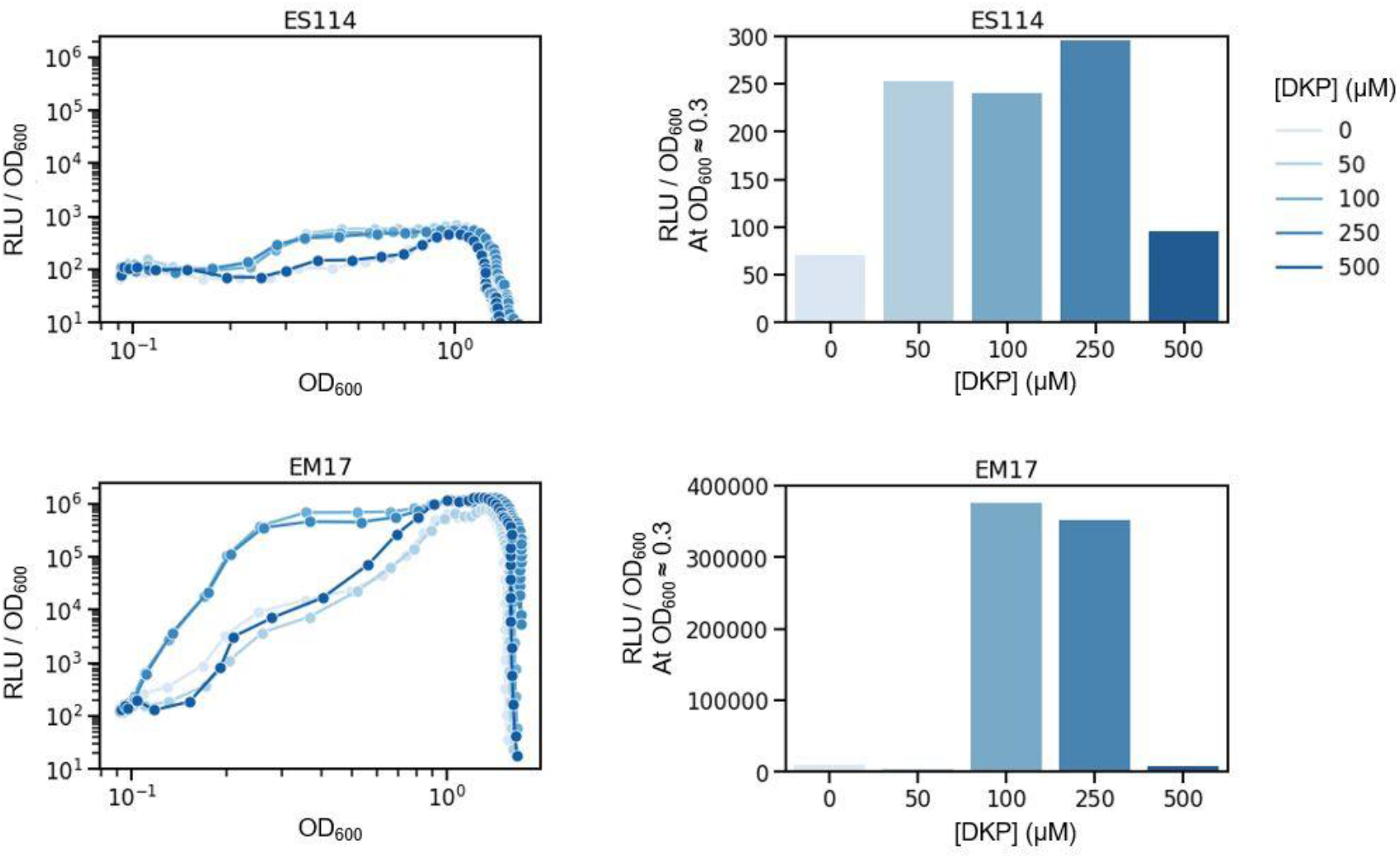
Cyclo(d-His-l-Pro) stimulates *V. fischeri* luminescence. Line graphs show cyclo(d-His-l-Pro) **3** increases relative light units in *V. fischeri* cultures at low concentrations in both a low-luminescence strain (ES114) and in a high-luminescence strain (EM17). Bar graphs show luminescence levels at a specific OD (OD600 ≈ 0.3) to illustrate the concentration-dependent effect. Graphs are representative of three independent experiments.

### Roles of Microbial Diketopiperazines

Microbial DKPs have been detailed in a number of biological contexts, including production in sexual pheromone signaling, as enzyme inhibitors, and in quorum sensing.(42, 63–65) Their structures vary widely, as there are many possible amino acid combinations and side chain modifications.(32) A thorough understanding of the ecology of the squid-*Vibrio* system provides context for the potential role of **3**. With an observable increase in production of **3** in a Biofilm-Up mutant, its possible to surmise that 3 may be critical for biofilm development, either as an end goal or as a preliminary step for a downstream process, or even as a quorum sensing molecule. Several bacterially derived DKPs have been described as chitinase inhibitors: cyclo(l-Arg-l-Pro), cyclo(Gly-l-Pro), cyclo(l-His,l-Pro) (**1**), and cyclo(l-Tyr,l-Pro).(65–67) While we have not explored the chitinase activity of **3**, if the stereoselectivity is not important in other activities of DKPs, it is possible that **3** displays the same inhibitory activity in the squid-*Vibrio* system, especially if **1** is known to be inhibitory. Because *E. scolopes* generates chitin as a source of nutrients for *V. fischeri*, presumably inhibition of chitinase by the bacterium would result in fewer monomers for consumption which would be thought to decrease fitness. Although this hypothesis has not yet been tested, there is precedent for the regulation of chitin and chitinase activity in this system.(68)

## Conclusion

The comparison of WT *V. fischeri* activity to mutants of strong and weak biofilm capabilities, *rscS* ΔbinK* and *rscS* rscS*::Tn*erm*, respectively, identified a small molecule, **3** that is produced in significant quantities in the Biofilm-Up (*rscS*\* *ΔbinK*) strain. **3** is a member of the DKP molecular class, members of which are increasingly ubiquitous across microbial species and have varying and dedicated activities including signaling, quorum sensing, and enzyme inhibition. The effect we observed on bacterial luminescence suggests that this DKP may link the early developmental process of biofilm formation with the later light production.

The biosynthetic origin of **3** is an immediate focus for our future work. Oftentimes, DKPs are shunt products to larger NRPS derived molecules, however, in our case, no larger NRPS has been detected in the media or in the genome, as of yet, which makes this biosynthetic route unlikely. Future studies will also focus on the protein target for luminescence activity, as well as the role in the light organ. The investigation of differentially produced small molecules in this controlled, two-partner system provides a platform for an increased understanding of the mechanisms that may be responsible for animal-microbe partners to recognize one another and to maintain a specific, life-long relationship.

## Acknowledgments

This publication was funded in part by the Chicago Biomedical Consortium with support from the Searle Funds at The Chicago Community Trust (LMS and MJM). Studies in the lab of LMS are supported by NIH grant R01GM125943-02S2 and UIC startup funds. Studies in the lab of M.J.M are supported by NIH grant R35GM119627 and NSF grant IOS-1757297. KEZ is supported by NIH grant F31 CA236237.

## Materials and Methods

### V. fischeri *Strains*

All strains are derivatives of the MJM1100 isolate of *V. fischeri* ES114, an *E. scolopes* light organ isolate (69, 70). MJM1776 is the “Biofilm-Down” isolate, with genotype MJM1100 *rscS* rscS*::Tn*erm*. It was isolated as a mariner transposon insertion from pMarVF1 in the *rscS* gene in strain MJM1198 (Mattias Gyllborg & M.J.M., unpublished) (71). MJM2255 is the “Biofilm-Up” isolate with genotype MJM1100 *rscS\** Δ*binK*, and which was described previously (25) MJM2251, with genotype MJM1100 Δ*binK* (25).

### Bioluminescence Assays

Two *Vibrio fischeri* strains, ES114 (WT) and *Euprymna morsei* symbiont EM17, were grown overnight in LBS at 25°C. Cultures were diluted 1/1000 in seawater tryptone with adjusted osmolarity (SWTO) with varying concentrations of **3** from 0 - 500 μM.(72) Samples were then transferred in triplicate to a Nunc clear bottom plate and measurements were taken by Biotek Synergy Neo2 plate reader. Relative luminescence (RLU) and OD600 were measured every 30 minutes for 22 hours. Triplicates were averaged and the specific luminescence (RLU/OD600) was plotted as a function of the OD600.

### Data Repository

Data files for all IMS and LS-MS/MS analysis can be found in the MassIVE Database under ID MSV000085327.

