## Supporting Information for "Discovery of a microbially produced small molecule in a host-specific organ"

General experimental and compound characterization………………………………. S2

**Figure S1.** Prioritized mass features detected using IMS…………………………. S12

**Figure S2.** GNPS match of *m/z* 235 to cyclo(His-Pro)....................................... S13

**Figure S3.** MALDI-TOF dried drop of m/z 235 in Biofilm-Up extract…………….. S14

**Figure S4.** Extracted ion chromatograms of cyclo(His-Pro) in Q-ToF…………... S15

**Table S1.** Optical rotation of all synthesized diastereomers……………………... S16

**Figure S5.** Retention time matching of cyclo(D-His-L-Pro) to Biofilm-Up……….. S17

**Figure S6.** Quantification of cyclo(D-His-L-Pro) in *V. fischeri* mutants………….. S18

References……………………………………………………………………………... S19
.

**General Experiment and Compound Characterization.**

V. fischeri *Growth and Plating for IMS*

Colonies on agar: Frozen stocks were revived by spreading onto an LBS agar (Luria-Bertani salt: 20 g NaCl, 10 g tryptone, 5 g yeast, 15 g agar, 50 mL 1 M Tris, pH 7.0 in 1 L DI $H_2O$) plate and after 24 h a single colony transferred to 5 mL LBS media.(1) Mutants were grown in liquid media for 24 h, and 5 µL of a normalized culture was spotted on 'thin' LBS agar plates (3 mL agar in 60 mm petri dishes). After 2 or 4 days of growth, *V. fischeri* colonies were excised on small patches of agar and transferred to a steel MALDI target plate. Colonies were covered with a layer of 50:50 dihydroxybenzoic acid (DHB): α-cyano-4-hydroxycinnamic acid (CHCA) using a 52 µm sieve and placed in a 37°C oven for at least 4 h or until dryness. Excess matrix was removed from the front and back of the plate using an air stream. A standard of phosphorus red (1 mg/µL, Sigma) was spotted onto a clear area of the plate and air dried.

*E. scolopes* hatchlings: Inoculation of *E. scolopes* hatchlings with *V. fischeri* mutants was performed as previously described in Zink *et al.* for the following mutants: No bacteria, MJM1100 (WT), and MJM2251 (WT ∆*binK*).(2)

*IMS Parameters and Data Analysis*

Colonies on agar: Regions of interest were designated using flexImaging v. 4.1 x 64 (Bruker) and data was acquired using flexControl v 3.4 at 200 µm spatial resolution on an Autoflex Speed LRF MALDI-TOF mass spectrometer (Bruker Scientific, Billerica, MA) over the mass range 100−1000 Da. In positive reflectron mode, laser power was set to 40%, laser width to 3 and reflector gain to 10×. For each raster point 500 laser shots at 500 Hz were shot in a random walk method. Data was subsequently analyzed in flexImaging v 4.1 × 64 (Bruker Scientific, Billerica, MA). All spectra were normalized to root mean square (RMS).

E. scolopes hatchlings: Hatchlings were thawed from -70 °C and prepared for IMS using the protocol described in Zink *et al*.(2) Light organ regions of interest (ROIs) were imaged at 20 μm spatial resolution with the following parameters: positive reflectron mode, laser power was set to 70%, laser width to 3 and reflector gain to 13×. For each raster point 500 laser shots at 2000 Hz were shot in a random walk method.

*Microbial Culture Growth and Extraction*

A frozen stock of Biofilm-Up was revived by spreading onto an LBS agar plate and after 24 h a single colony transferred to 5 mL LBS media. 500 mL LBS agar was solidified in a 12 x 8 in autoclave tub and the 5 mL liquid culture spread evenly across the surface of the agar. Once dried, the tub was covered with cheesecloth and secured with rubber bands, then flipped upside down and left for 4 days at 25°C. After four days, the top layer of agar along with the biofilm was collected using a spatula and submerged in 200 mL of 1:1 $H_2O$:MeOH. After 1 h of sonication, the agar and biofilm mass was filtered out from the solvent using a cheesecloth filter. 8 g of 'polar' resin (1:1:1: of XAD2:XAD4:XAD7 (Sigma)) were added to the solvent and shaken at 225 rpm for 1 h, then the resin was filtered out using a cheesecloth filter and back-extracted in 100 mL of 1:1 ACN:$H_2O$ + 0.1% TFA, shaking at 225 rpm for 1 h. After 1 h, the back extract was filtered from the resin using a cheesecloth filter and dried under rotoevaporation. Dried extractions were fractionated by polarity using solid phase extraction (Discovery 5 g SPE column) over nine increments: 5% MeOH, 10% MeOH, 15% MeOH, 20% MeOH, 40% MeOH, 60% MeOH, 80% MeOH, 100% MeOH, and 100% EtOAc. Each fraction was dried *in vacuo* and analyzed via dried drop on the MALDI-TOF MS.

*GNPS Parameters*

Crude extracts of WT, Biofilm-Down, and Biofilm-Up were diluted to 1 mg/mL in MeOH and analyzed on a Thermo Finnigan LCQ Advantage Max under the following parameters. Raw data files were converted to mzXML using MSConvert and uploaded to gnps.ucsd.edu for molecular networking. Controls for the experiment included an LBS extract and a MeOH blank. Molecular networking was analyzed under the following parameters: Precursor Ion Mass Tolerance: 2.0 Da, Fragment Ion Mass Tolerance: 0.5 Da, Min Pairs Cos: 0.6 Da, Minimum Matched Fragment Ions: 3.

The crude extract of Biofilm-Up and a 500 nM solution of a standard of **3** was run on a Compact II Q-TOF (Bruker) to obtain a high resolution mass with the following parameters: LC 2-10% B over 10 min (A: $H_2O$ + 0.2% FA, B: ACN + 0.2% FA), positive mode, 100-2000 Da, Top 9 precursor ions fragmented at 35 eV per scan. An ESI TOF mix (Bruker) was used as a calibrant between 7.8 to 8.0 min. Fragmentation using direct infusion was acquired by using a crude extract of Biofilm-Up compared to a 10 mM solution of **1**.

*Compound Characterization*

Three stereoisomers (**2** - **4**) were synthesized following the protocol outlined in Kukla *et al.(3)* Final products were dried under high vacuum to give either a white powder (**3**) or a yellow oil (**2**, **4**). Stereoisomer **1** was purchased from Sigma. For easy reference, the protocol used to prepare **3** is given below; stereoisomers **2** and **4** were prepared using analogous procedures.

Hydroxybenzotriazole (HOBt, 60 mg, 0.1 eq.), ((benzyloxy)carbonyl)-L-proline (Cbz-L-Pro, 1.00 g, 1.0 eq.), (1-[Bis(dimethylamino)methylene]-1H-1,2,3-triazolo- [4,5-b]pyridinium 3-oxid hexafluoro-phosphate) (HATU coupling reagent, 1.54 g, 1.0 eq.) were combined in a two-neck round-bottom flask, secured above a stir plate and equipped with a stir bar. Dichloromethane (22.5 mL) and *N,N*-diisopropylethylamine (3 mL, 4.3 eq.), were added to the flask, and the mixture was stirred for 2-3 min. D-Histidine methyl ester dihydrochloride (980 mg, 1.0 eq.)was added to the flask. The solution was stirred for 2.5-3 h, and the reaction progress was monitored by HPLC.

$H_2O$ (ca. 150 mL) was added, and the reaction mixture was transferred to a separatory funnel. The reaction mixture was extracted twice with dichloromethane. The combined organic extracts were dried over $MgCl_2$. The filtrate was reduced to dryness on a rotary evaporator. The crude product was purified using automated flash column chromatography (40 g silica column, 0-5% 90:10 DCM:MeOH.) The product eluted at 2.5%. Fractions containing the product, as determined by thin layer chromatography, were combined and evaporated under rotary evaporation and high vacuum to give 1.30 g of Cbz-L-Pro-D-His-OMe (81% yield).

Purified Cbz-L-Pro-D-His-OMe (1.199 g, 1.0 eq.) was transferred to a hydrogenation vessel to which palladium on carbon (256 mg of 50% wet, 10 wt. % Pd) was added. The reaction vessel was evacuated under vacuum for 3 min. and was backfilled with argon three times. MeOH (23 mL) was added to the flask under argon. The vessel was transferred to a Parr Shaker apparatus and filled and backfilled three times with $H_2$ gas (~40 psi). The vessel was shaken vigorously for 1 hr. The product was vacuum-filtered through celite with the aid of MeOH (2X) to remove Pd. The filtrate was dried under rotovap and left under high vacuum to dryness to give 790 mg of H-L-Pro-D-His-OMe (99% yield).

Cyclization was promoted by dissolving 790 mg H-L-Pro-D-His-OMe powder in 50 mL MeOH in a round bottom flask equipped with a stir bar and condenser. The reaction mixture was stirred at reflux overnight. After the reaction was completed, as determined by LC-MS, the solvent was removed via rotary evaporation and dried under high vacuum. The crude products were purified by semi-preparative HPLC, as described below. Purity of cyclo(D-His-L-Pro) was checked on HPLC.

*Isolation of final products*

Small-scale synthesis products were isolated from byproducts using semi-preparative HPLC under the following conditions: 1% B isocratic over 15 min (A: DI $H_2O$ + 0.1% TFA, B: ACN + 0.1% TFA) on C-18 Kinetex column (Phenomenex, 2.6 µm PS C18 100 Å, LC Column 50 x 4.6

mm). UV monitoring at 210 nm, and products collected at 8.0-9.0 min. Products dried under high vacuum.

Large scale synthesis of **3** was isolated by combiflash under the following conditions: 40 g silica column, 0-5% 90:10 DCM: MeOH. **3** collected at ~2.5 min and dried under high vacuum.

All products were verified for stereochemistry using previously reported NMR and optical rotation values (**Table S1**).(3)

*HPLC Retention Time Matching*

Two 1:1 racemic mixtures (**1** and **4**, **2** and **3**) along with a 1:1:1:1 mixture of **1-4** were sent to Regis Technologies, Inc. for chiral screening. The best separation of all four stereoisomers was observed on the Reflect C-Cellulose B column, 250 mm x 4.6 mm, particle size 5 µm. Screening conditions were adapted in-house to the following for retention time matching: 13% B isocratic over 60 min (A: Hexanes, B: IPA), flow rate = 2.0 mL/min. An Agilent Technologies 1260 Infinity HPLC equipped with DAD detector was used. All solvents used were Optima grade and were not further purified.

*Quantification of **3** in* V. fischeri *Cultures*

Earlier evaluation of several different growth types assessed that growth of *V. fischeri ΔbinK* on large-scale agar as solid colonies generated the highest amount of **3** for extraction. In attempts to quantify the DKP from culture, all mutants were grown at several time points and both in liquid and solid media. Small-scale samples (5 mL in liquid, 100 µL inoculation on agar) were grown and extracted after three individual time points in triplicate: 24, 48, and 96 hours. Extracts were analyzed on a high-resolution mass spectrometer and the biofilm extracts after 48 hours of growth on agar indicated that *ΔbinK* produced a significantly higher abundance of **3** than both the

WT and the disadvantaged colonizer, *rscS* (**Figure S6**). T-tests calculated the significant difference of production with both mutants individually to *ΔbinK* is $p < 0.05$ (*).

The consistency in the significant production of the DKP at the 48 h time-point *in vivo* and *in vitro* indicates that this period of inoculation may be biologically relevant.

*Quantitation of **3** via LC-MS/MS*

Biofilm-Down, WT, and Biofilm-Up were grown in triplicate at three time points (24 h, 48 h, and 96 h) in both liquid culture and in biofilms on agar plates. Liquid cultures were grown as follows: bacteria were inoculated in 5 mL of LBS from frozen stocks spread on agar plates and incubated overnight. Bacteria were diluted 1/100 in 5 mL LBS in triplicate and incubated at 37°C, shaking at 225 rpm. After 24, 48, or 96 h, 1 g of 'polar' resin (XAD 2:4:7) was added to the cultures and shaken at 225 rpm and 37°C for 1 h. Resin was vacuum-filtered out of the culture and back-extracted using 50:50 $H_2O$:MeOH + 0.1% TFA, shaking at 225 rpm and 37°C for 1 h. Extractions were dried *in vacuo* at 30 °C and resuspended in DI $H_2O$ at 1 mg/mL for Q-TOF analysis.

Biofilm cultures were grown as follows: bacteria were inoculated in 5 mL of LBS from frozen stocks spread on agar plates and incubated overnight. 100 µL of liquid culture was spotted onto agar LBS plates in triplicate. After 24, 48, or 96 h, LBS was removed from the test tube using an automated pipetman, and 5 mL of DI $H_2O$ was added, then shaken at 225 rpm and 37 °C for 1 h. Resin was vacuum-filtered out of the culture and back-extracted using 50:50 $H_2O$:MeOH + 0.1% TFA, shaking at 225 rpm and 37°C for 1 h. Samples were centrifuged at 10k rpm for 2 min to concentrate remaining biofilm material at the bottom of the tube. Extractions were dried *in vacuo* at 30°C and resuspended in DI $H_2O$ at 1 mg/mL for Q-TOF analysis.

LC analysis utilized 2-10% B over 10 min (A: $H_2O$ + 0.2% FA, B: ACN + 0.2% FA), and MS parameters utilized positive mode, 100-2000 Da, Top 9 precursor ions fragmented at 35 eV per scan. An ESI TOF mix (Bruker) was used as a calibrant between 7.8 to 8.0 min. The values of intensity were generated in Metaboscape.

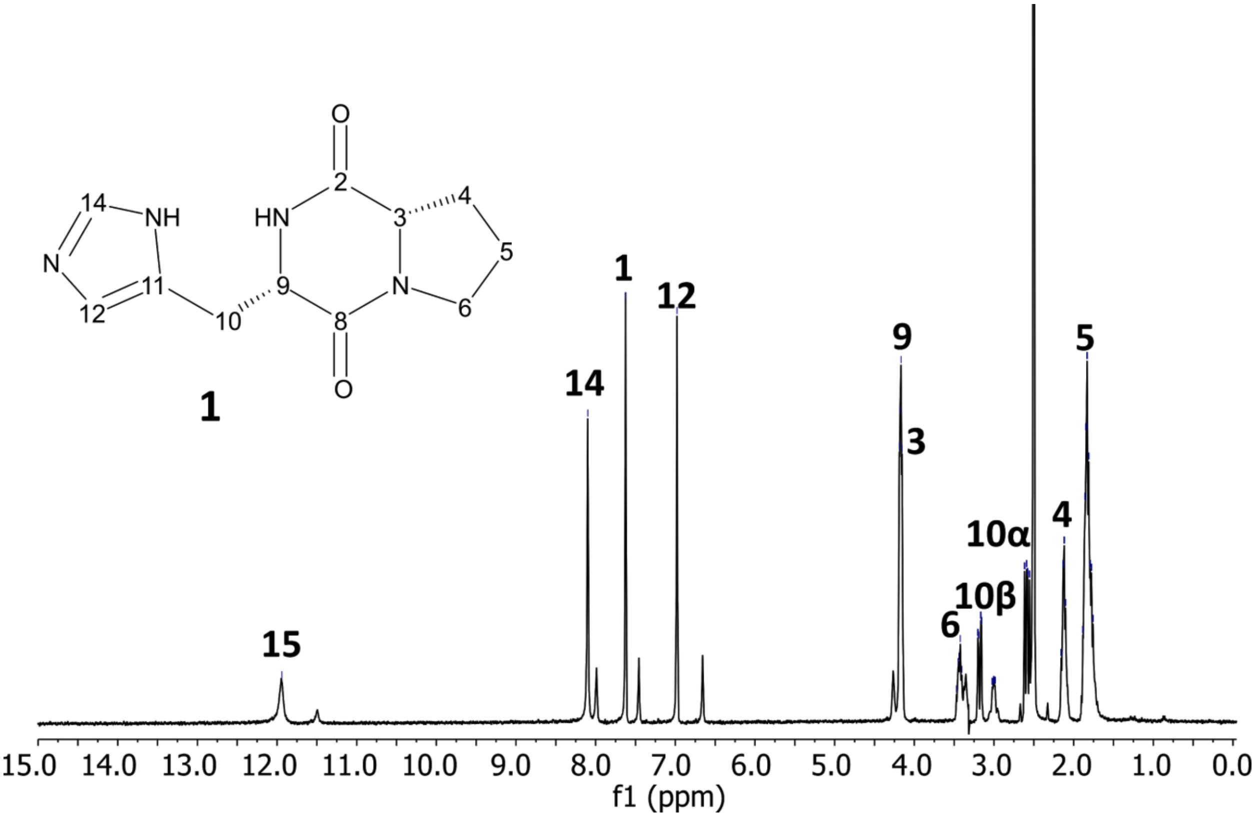


Cyclo(L-His-L-Pro) **1**: White solid. $[\alpha]_D^{20}$ -132 (*c* 0.001, MeOH). UV ($H_2O$) $\lambda_{max}$ (ε log) 214 nm. $^1$H NMR (400 MHz, DMSO-$d_6$) δ 11.94 (s, 1H), 8.10 (s, 1H), 7.62 (s, 1H), 6.98 (s, 1H), 4.17 (dd, *J* = 9.3, 4.3 Hz, 2H), 3.49 – 3.31 (m, 1H), 3.18 (dd, *J* = 14.9, 3.4 Hz, 1H), 3.09 – 2.92 (m, 1H), 2.58 (dd, *J* = 14.9, 9.7 Hz, 1H), 2.15 – 2.08 (m, 1H), 1.88 – 1.72 (m, 4H). HRESIMS *m/z* $[M+H]^+$ 235.1190 (calcd for $C_{11}H_{15}N_4O_2$, 235.1195).

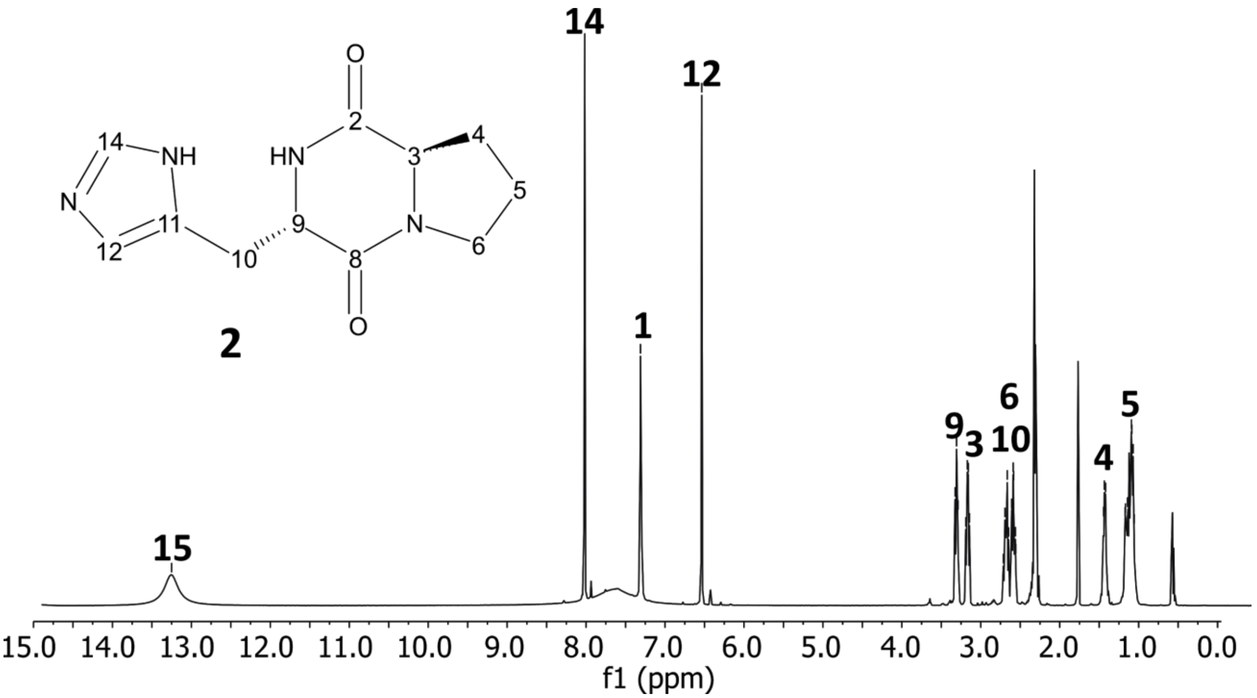


Cyclo(L-His-D-Pro) **2**: Yellow oil. $[\alpha]_D^{20}$ +12 (*c* 0.001, MeOH). UV ($H_2O$) $\lambda_{max}$ (ε log) 214 nm. $^1H$ NMR (400 MHz, DMSO-$d_6$) δ 14.47 (s, 1H), 9.02 (s, 1H), 8.28 (s, 1H), 7.48 (s, 1H), 4.11 (t, J = 7.8 Hz, 1H), 4.00 – 3.93 (m, 1H), 3.50 – 3.40 (m, 1H), 3.41 – 3.31 (m, 1H), 3.11 – 3.04 (m, 2H), 2.20 – 2.11 (m, 1H), 1.84 (ddd, J = 26.8, 10.4, 3.8 Hz, 3H). HRESIMS *m/z* $[M+H]^+$ 235.1190 (calcd for $C_{11}H_{15}N_4O_2$, 235.1195).

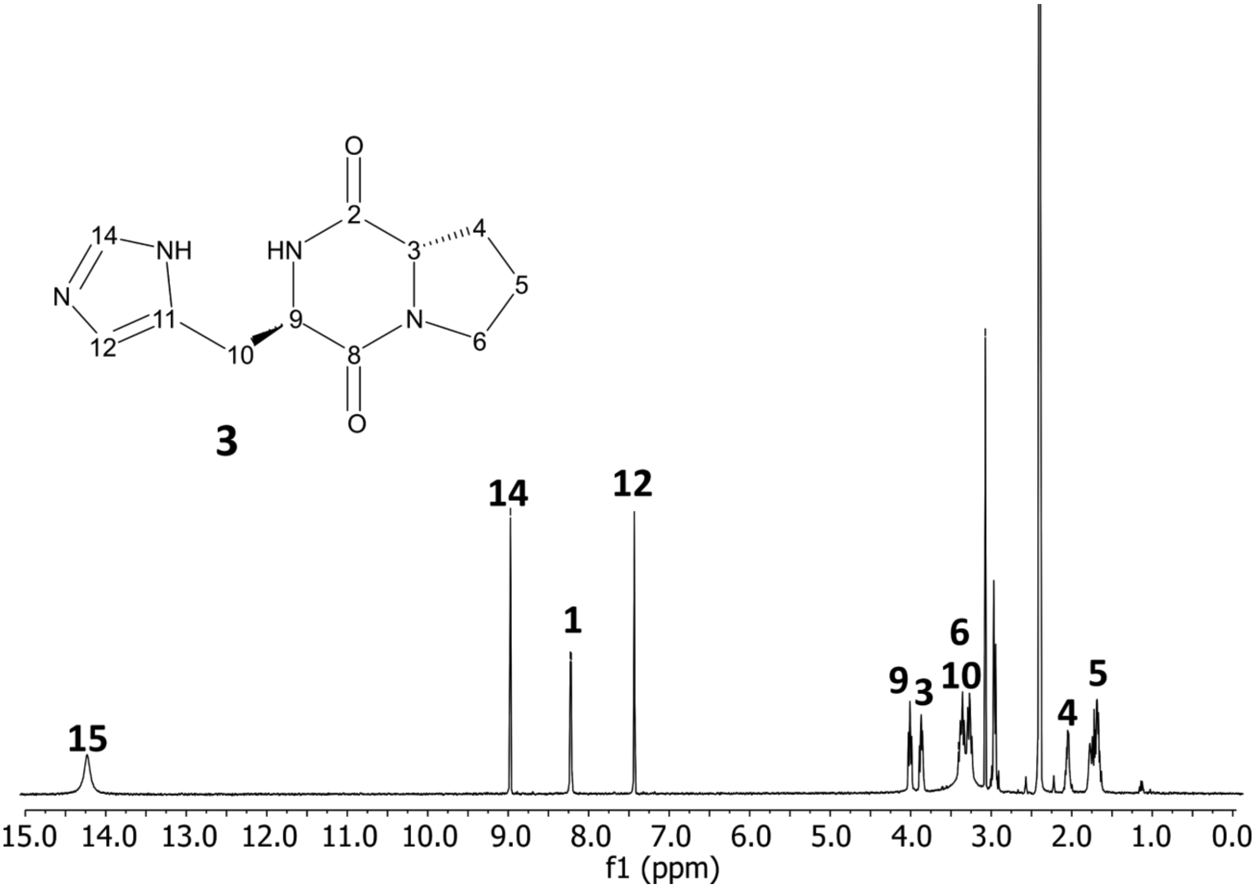


Cyclo(D-His-L-Pro) **3**: White solid. $[\alpha]_D^{20}$ -7 (*c* 0.001, MeOH). UV ($H_2O$) $\lambda_{max}$ (ε log) 214 nm. $^1H$ NMR (400 MHz, DMSO-$d_6$) δ 14.17 (s, 1H), 8.99 (s, 1H), 8.25 (d, J = 4.2 Hz, 1H), 7.47 (s, 1H), 4.09 (t, J = 7.8 Hz, 1H), 3.98 – 3.91 (m, 1H), 3.51 – 3.41 (m, 2H), 3.35 (ddd, J = 11.0, 8.3, 2.4 Hz, 2H), 3.17 (s, 1H), 3.06 (s, 1H), 3.05 (d, J = 4.3 Hz, 1H), 2.19 – 2.12 (m, 1H), 1.92 – 1.74 (m, 3H). HRESIMS *m/z* $[M+H]^+$ 235.1190 (calcd for $C_{11}H_{15}N_4O_2$, 235.1195).

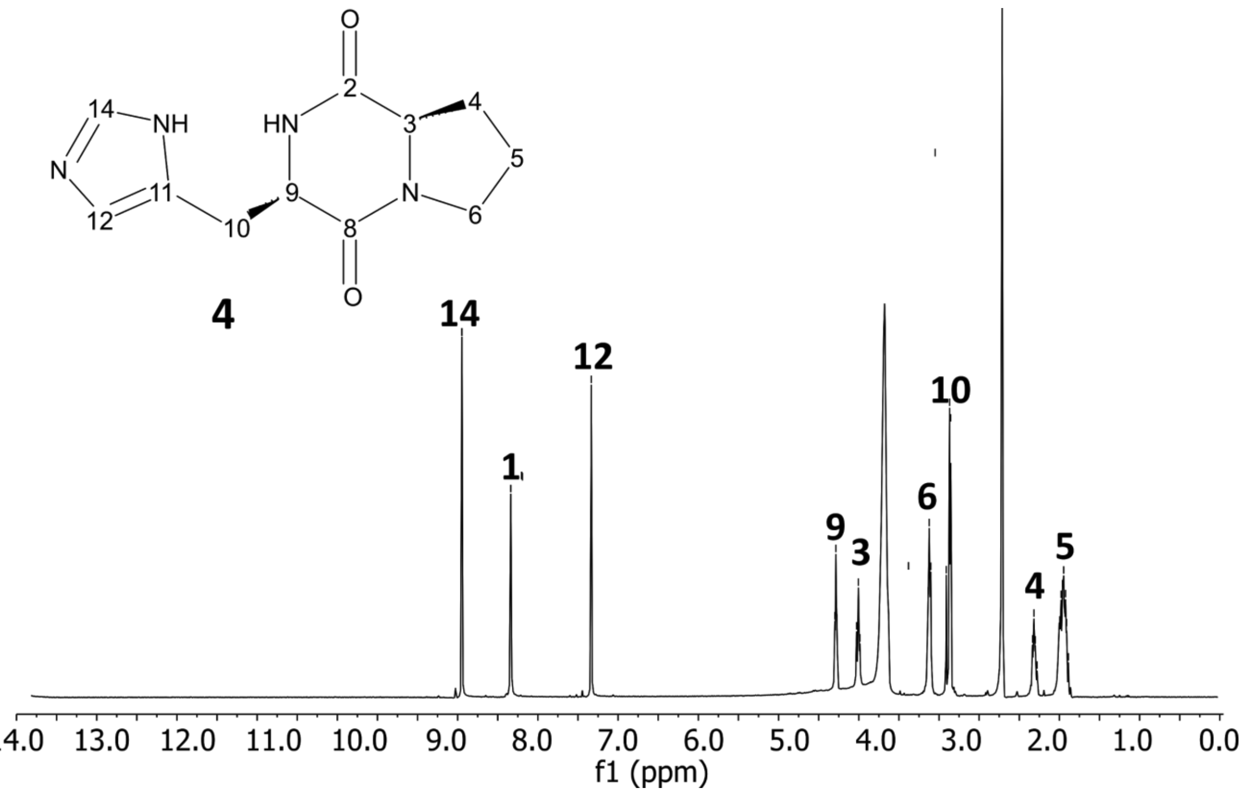


Cyclo(D-His-D-Pro) **4**: Yellow oil. $[\alpha]_D^{20}$ +47 (*c* 0.001, MeOH). UV ($H_2O$) $\lambda_{max}$ (ε log) 214 nm. $^1$H NMR (400 MHz, DMSO-$d_6$) δ 8.79 (s, 1H), 8.22 (s, 1H), 7.28 (s, 1H), 4.44 (t, *J* = 4.9 Hz, 1H), 4.18 (t, *J* = 7.6 Hz, 1H), 3.38 – 3.31 (m, 2H), 3.11 (d, *J* = 5.2 Hz, 2H), 2.16 – 2.08 (m, 1H), 1.78 (tt, *J* = 13.4, 7.2 Hz, 3H). HRESIMS *m/z* $[M+H]^+$ 235.1190 (calcd for $C_{11}H_{15}N_4O_2$, 235.1195).

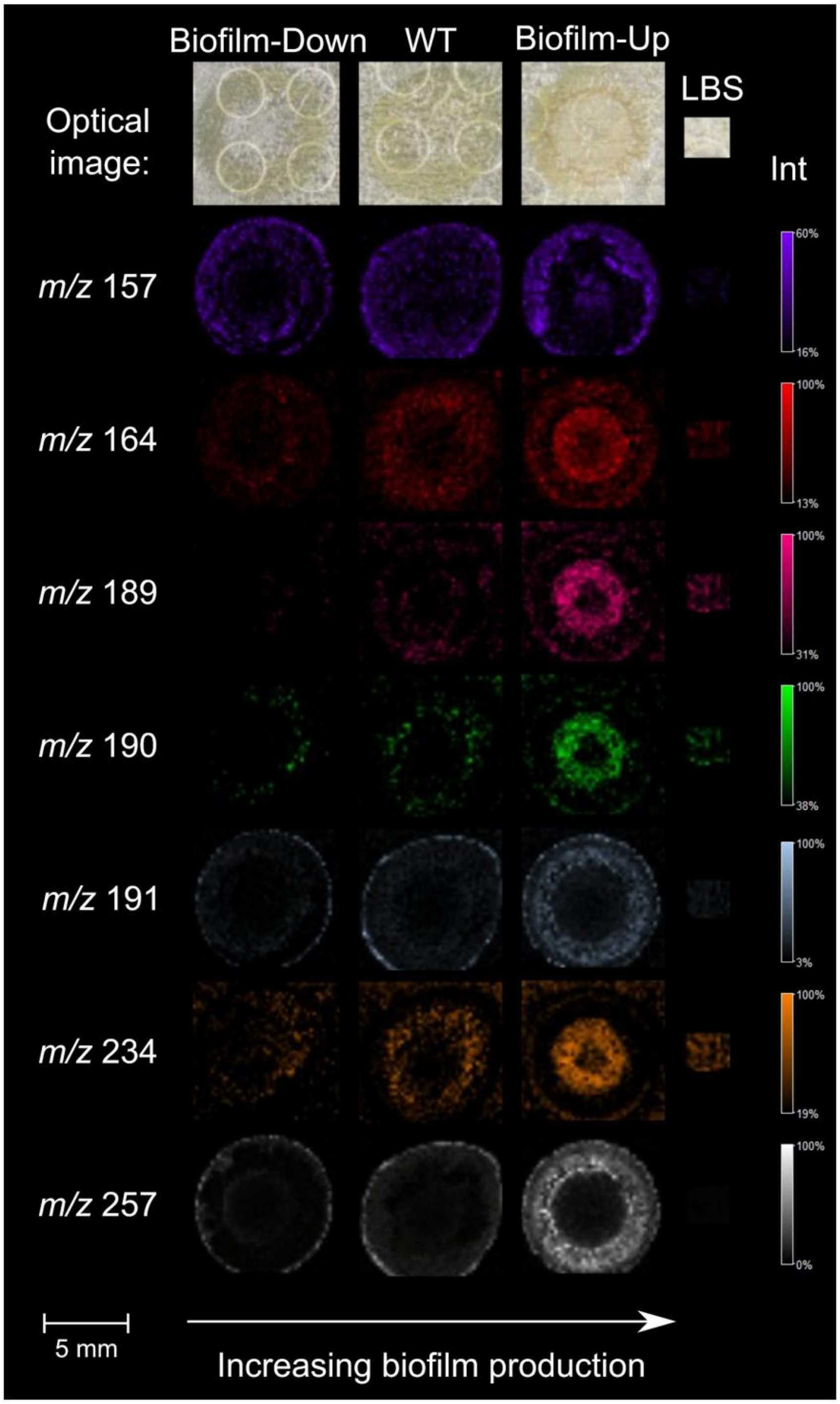


**Figure S1.** Seven small molecules were significantly more abundant ($p < 0.1$) in Biofilm-Up compared to WT and Biofilm-Down, detected as significant at least two across four biological replicates. IMS analysis was performed in positive mode and in the mass range of 100-1000 Da.

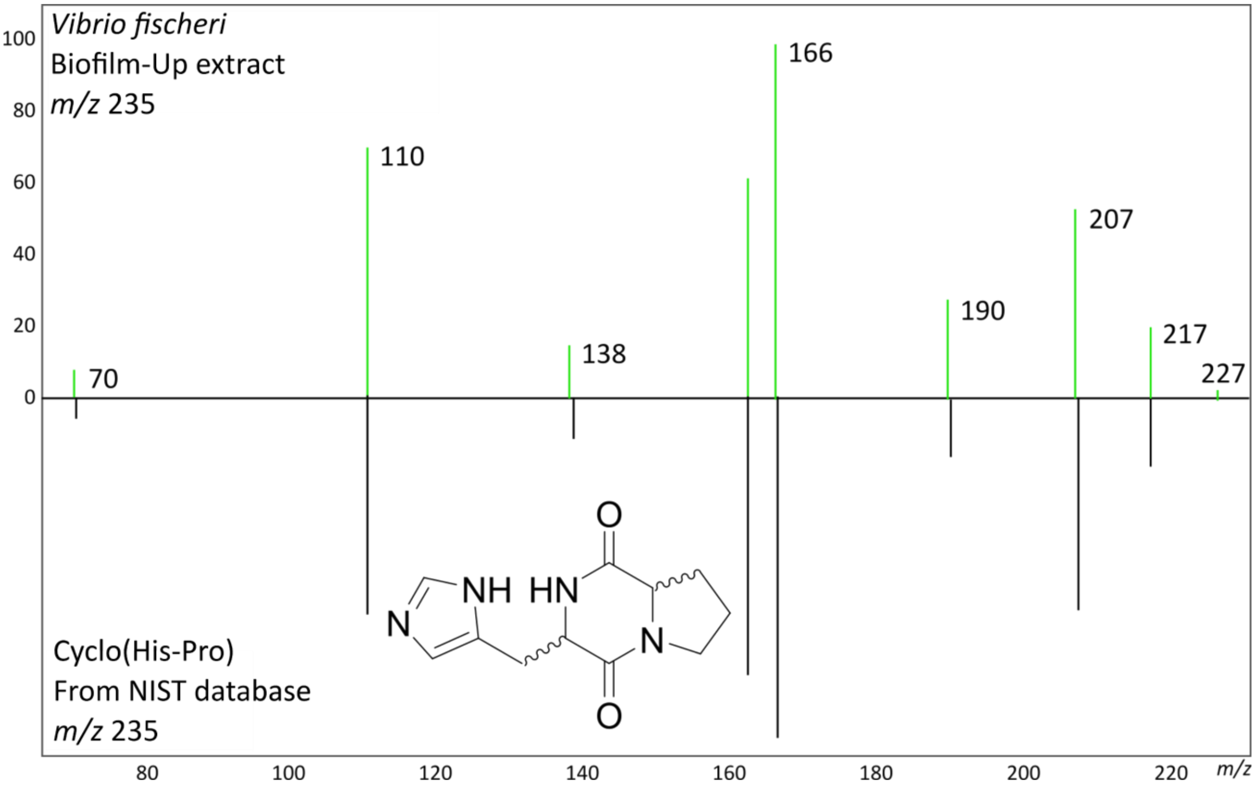


**Figure S2.** The crude extract of *V. fischeri* Biofilm-Up was queried in the GNPS database and a strong match was calculated between a molecule in the crude extract and cyclo(histidyl-proline), *m/z* 235. The compound was added from the NIST database under the Library Spectrum CCMSLIB00003139663.

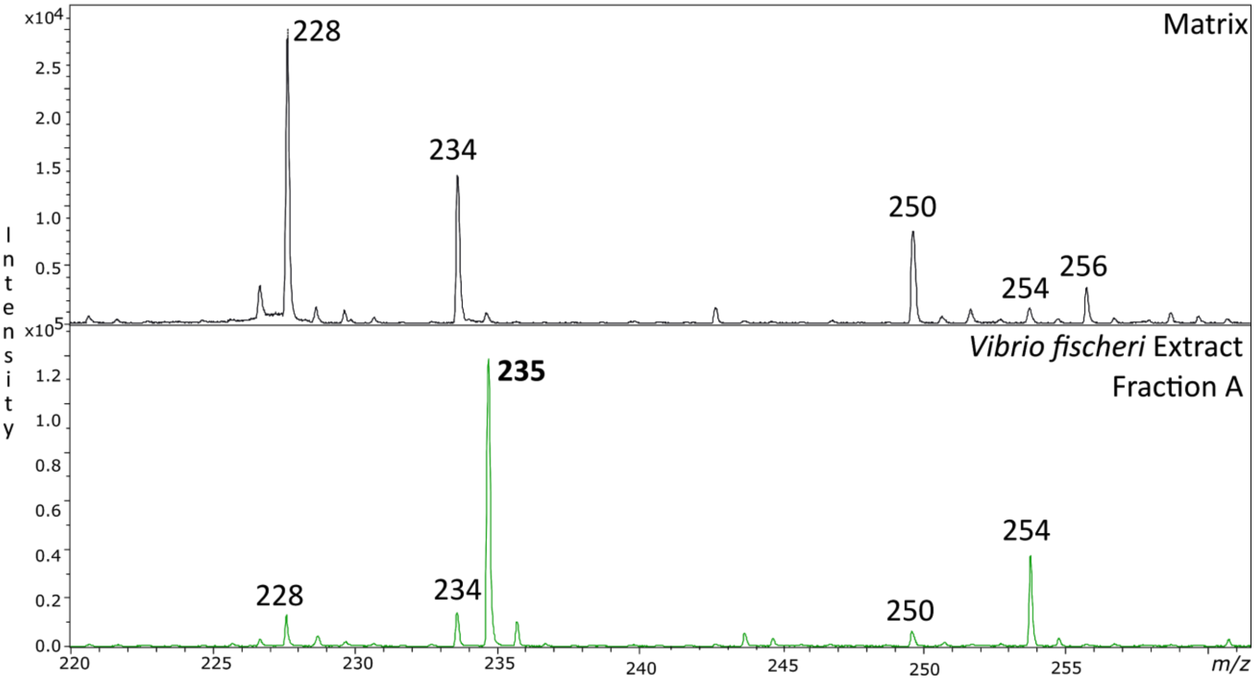


**Figure S3.** Dried drop analysis of Fraction A from *V. fischeri* Biofilm-Up extract. The *m/z* 235 is present in the extract fraction, but not in the matrix control.

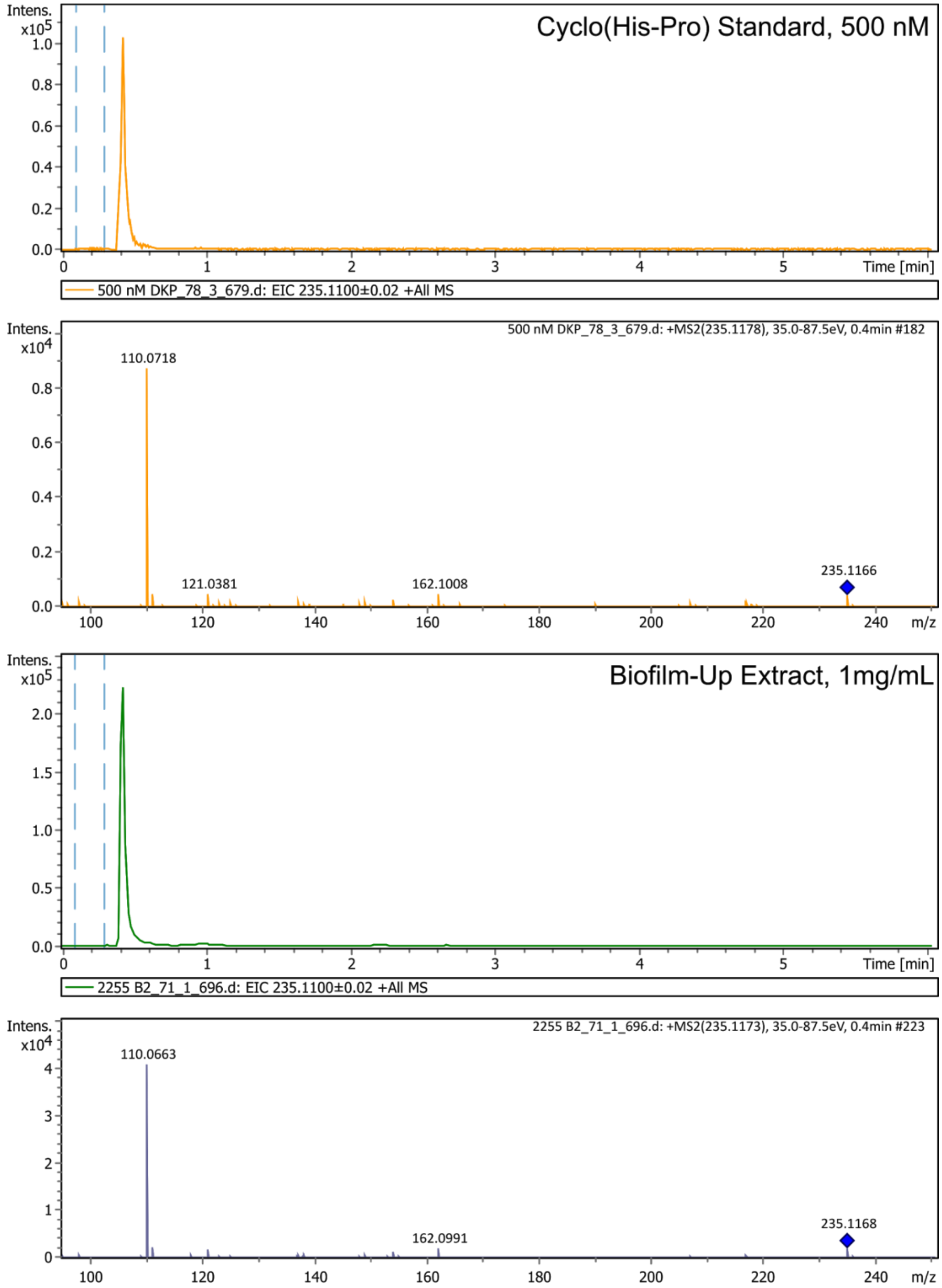

**Figure S4.** High-resolution electrospray ionization mass spectrometry (HRESIMS) of Cyclo(His-Pro) compared to the extract of Biofilm-Up *Vibrio fischeri* strain. Extracted ion chromatograms (EICs) of *m/z* 235.11 +/- 0.2 Da detected a precursor ion of 235.1178 in the standard, and 235.1173 in the extract (ppm error 7.23 and 6.80, respectively).

| Configuration | $[\alpha]_D$ experimental | *c* experimental | $[\alpha]_D$ reported | *c* reported |
|---|---|---|---|---|
| Cyclo(L-Histidine,L-Proline) **1** | -132 | 0.001 | -119 | 1.0 |
| Cyclo(L-Histidine,D-Proline) **2** | +12 | 0.001 | +123 | 0.96 |
| Cyclo(D-Histidine,L-Proline) **3** | -7 | 0.001 | -53.8 | 0.31 |
| Cyclo(D-Histidine,D-Proline) **4** | +47 | 0.001 | +53.8 | 0.94 |

**Table S1.** Optical rotation values for all four diastereomers of cyclo(histidyl-proline) compared to literature values from paper that outlined synthesis procedure. All compounds, reported and experimental, were measured in MeOH. Concentrations (*c,* g/mL) of compounds synthesized in this work are 1000 times less than those reported by Kukla *et al*, which may be responsible for discrepancies in specific rotation values.(3)

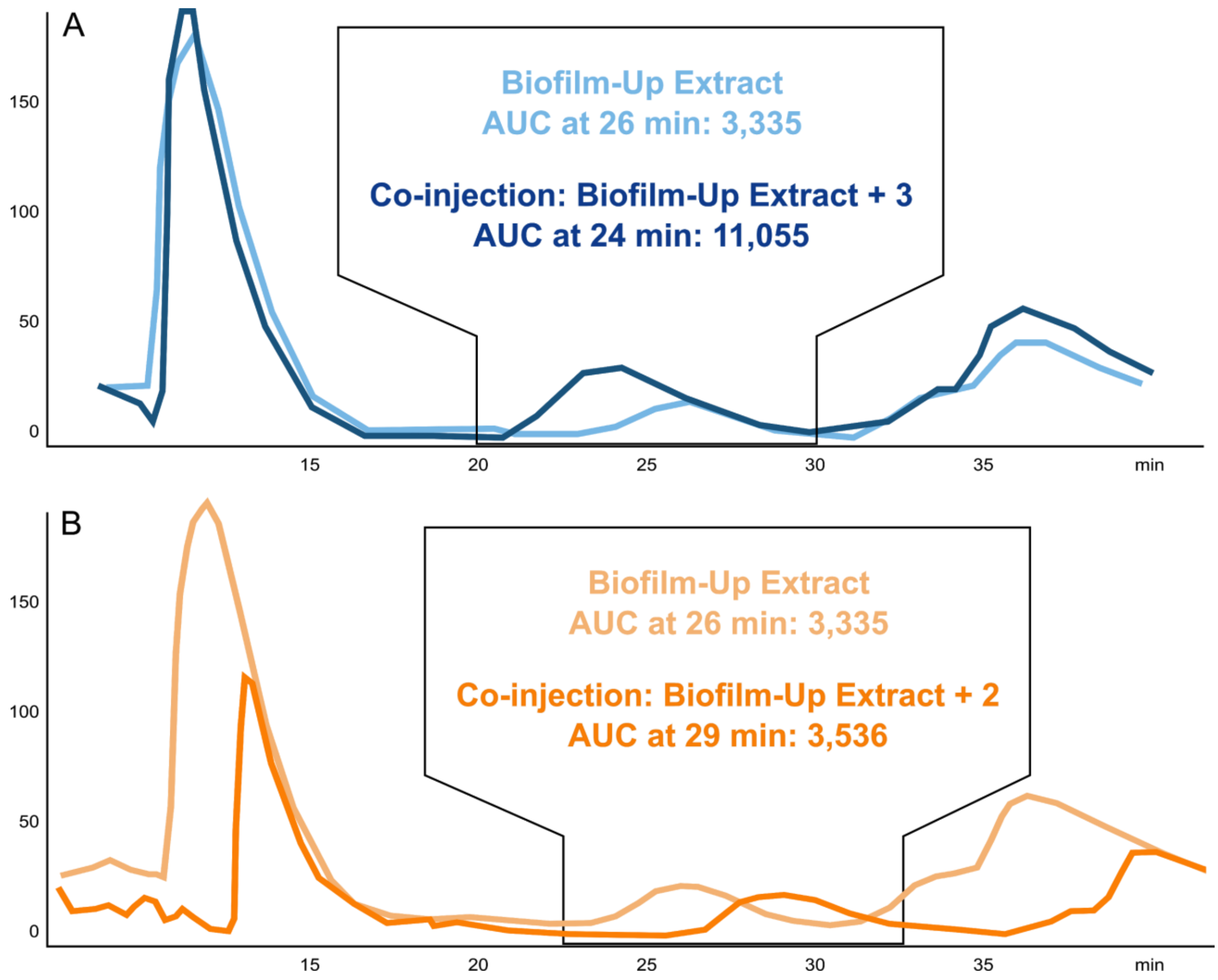


**Figure S5**. Retention time (RT) matching of stereoisomers compared to Biofilm-Up extract. A peak was observed in the extract at 26 min with an area under the curve (AUC, indicative of the peak intensity) of 3,335. Cyclo(D-His-L-Pro) (**3**) and cyclo(L-His-D-Pro) (**2**) eluted from the chiral column at RTs of 24 min and 29 min, respectively, indicating that the peak from the extract at 26 may represent one of the two stereoisomers. **A)** Co-injection of the extract with **3** resulted in elution of a peak at 24 min whose AUC increased to 11,055. **B)** Co-injection of the extract with **2** resulted in a peak at 29 min with an AUC of 3,536, indicating that the extract peak and DKP peak did not coalesce, and therefore do not share the same configuration.

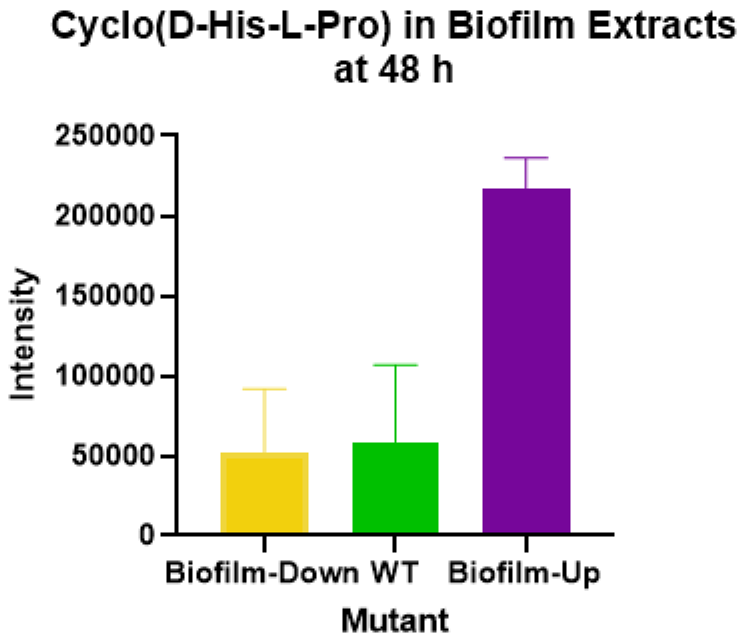


**Figure S6.** Quantification of **3** *in vitro* in *V. fischeri* mutants demonstrated that the Biofilm-Up strain produced significantly more of **3** than the WT and the Biofilm-Down strain.
